# Propionic acid promotes the virulent phenotype of Crohn’s disease-associated adherent-invasive *Escherichia coli*

**DOI:** 10.1101/387647

**Authors:** Michael J. Ormsby, Síle A. Johnson, Lynsey M. Meikle, Robert J. Goldstone, Anne McIntosh, Hannah M. Wessel, Heather E. Hulme, Ceilidh C. McConnachie, James P. R. Connolly, Andrew J. Roe, Conor Hasson, Joseph Boyd, Eamonn Fitzgerald, Konstantinos Gerasimidis, Douglas Morrison, Georgina Hold, Richard Hansen, David G.E. Smith, Daniel M. Wall

**Author notes:** Corresponding author address: Dr. Daniel (Dónal) M. Wall, Institute of Infection, Immunity and Inflammation College of Medical, Veterinary and Life Sciences, Sir Graeme Davies Building, University of Glasgow, 120 University Place, Glasgow G12 8TA, Corresponding author.

## Abstract

The short chain fatty acid propionic acid (PA) is a bacteria-derived human intestinal antimicrobial and immune modulator used widely in Western food production and agriculture. Here we examine the effect of PA on the pathogenicity of the Crohn’s disease-associated microbe, adherent-invasive *Escherichia coli* (AIEC). Passage of AIEC through a murine model, where the low intestinal PA levels were increased to replicate those of the human intestine, led to the recovery of AIEC post-infection that had significantly increased virulence. These phenotypic changes, including increased adhesion to intestinal epithelial cells and biofilm formation, could be replicated in AIEC *in vitro* through exposure to PA alone. This *in vitro* exposure of AIEC to PA fundamentally changed AIEC virulence, with strains exposed to PA *in vitro* subsequently persisting at 20-fold higher levels in a murine model compared to non-exposed strains. RNA-sequencing identified the transcriptional changes in AIEC in response to PA with upregulation of genes involved in biofilm formation, stress responses, metabolism, membrane integrity and alternative carbon source utilisation. These PA induced changes in virulence could be replicated in a number of *E. coli* isolates from Crohn’s disease patients. Finally, removal of the PA selective pressure was sufficient to reverse these phenotypic changes. Our data indicate that exposure of AIEC to PA evolves bacteria that are both resistant to this natural human intestinal antimicrobial and increasingly virulent in its presence.

**Importance:** Exposure to propionic acid, an intestinal short chain fatty acid and commonly used antimicrobial in Western food production, induces significant virulence associated phenotypic changes in adherent-invasive *Escherichia coli* (AIEC).

## Introduction

Short chain fatty acids (SCFAs) are naturally produced by gut bacteria, through the breakdown of undigested carbohydrates and starches. This process results in the production of acetic (AA), butyric (BA) and propionic acids (PA) which accounts for approximately 90% of intestinal SCFAs. PA has attracted significant interest due to its potent immunomodulatory effects, with its supplementation shown to reduce the severity of the colitis in murine models, suggesting its modulation has potential as a therapeutic intervention in inflammatory bowel disease *(1, 2)*. Crohn’s disease (CD) is a debilitating and incurable inflammatory disease of a multi-factorial aetiology. The mechanisms underlying the disease are not fully understood, however it is thought that defects in the immune response to the gut microbiota are a contributing factor *(3–8)*. Sudden changes in diet have been shown to result in rapid changes in the gut microbiota *(9–11)*, while the inflammation associated with CD results in markedly decreased microbial diversity *(12–14)*. Levels of *Enterobacteriaceae* in particular are higher in intestinal samples from CD patients when compared to healthy controls *(15–17, 7)*. One group of *Enterobacteriaceae* that is of particular interest is the *Escherichia coli* pathotype, adherent-invasive *E. coli* (AIEC). These bacteria are overrepresented in the ileal microbiota of CD patients, being present in 51.9% of mucosal samples from CD patients compared with 16.7% in healthy controls *(18)*. Key features of the AIEC pathotype that distinguish them from non-invasive commensal strains include; adherence to and invasion of the intestinal epithelium, an increased ability to form biofilms, and the ability to survive and replicate within macrophages without inducing cell death *(18)*. While AIEC strains harbour genetic similarity to extra-intestinal pathogenic *E. coli* (ExPEC) in terms of phylogenetic origin and virulence genotype, they have few known virulence factors *(19)*. This apparent lack of virulence factors and the discovery of AIEC strains across all five major diverse phylogroups of *E. coli*, mean that an overarching explanation for the origin and virulence of AIEC has remained out of reach.

PA is manufactured on an industrial scale and is now commonly used in agriculture as in addition to its anti-inflammatory effects it is a potent antimicrobial. PA has demonstrated success in reducing pathogen numbers in poultry, particularly in reducing *Salmonella* and *Campylobacter* carriage *(20–23)*. PA is also an effective antimicrobial agent used in Western food production and agriculture, reducing the need for antibiotic use amidst growing antibiotic resistance concerns *(24–26)*. Inclusion in animal feed, grain, and food for human consumption accounted for almost 80% of PA consumption across the world in 2016, with Western Europe (40% of total use), North America (30%) and Asia (23%), the main consumers *(27)*. The success of SCFAs in reducing antibiotic dependence is now seeing their use spread to countries across Africa, the Middle East and Central and South America *(27)*. While there is increasing evidence for antibiotic-driven enhanced genome wide mutation rates and horizontal transmission of bacteria from food producing animals to humans *(28–31)*, the role of alternative antimicrobials such as PA in such phenomena has yet to be addressed *(32–34)*. Indeed, using animal models for human disease where SCFAs may play an important role has proved difficult at best. Murine models for human pathogens are limited by distinct differences in basal levels of SCFAs between the murine and human intestines with murine levels significantly lower in the case of PA *(35)*. However the significance of such differences in influencing the outcome or course of disease is not known but could be substantial.

In this study, we show that PA-exposure promotes increased virulence in AIEC. Using a murine model with increased intestinal PA concentrations, we have generated a more relevant model for human gut-associated AIEC infection, resulting in increased AIEC virulence and persistence. This increased virulence is PA-dependent and can be replicated *in vitro* by exposure of AIEC to PA alone. RNA sequencing identified the transcriptional changes driving these changes in virulence, which were determined to be reversible if PA was removed. Our data highlights the potential risks of widespread PA use as an antimicrobial. An understanding of the AIEC phenotype using genetic markers or phenotypic characteristics has remained elusive, however, our findings indicate that PA metabolism is a crucial driver of the AIEC phenotype.

## Results

### Humanising the murine intestinal PA concentration exacerbates the AIEC phenotype

While the concentration of PA in the human intestine ranges from 1.5 mM in the ileum to 27 mM in the colon, it is considerably lower in the murine intestine (Fig. 1a) *(20, 35)*. To address this in our *in vivo* infection model, we supplemented the drinking water of male C57BL6 mice with 20 mM PA. Caecal SCFA levels post-PA supplementation indicated that PA concentration had significantly increased while no changes were seen in either AA or BA concentrations (Supplementary Fig. S1; AA:PA:BA ratios without PA 84.6:7.3:8.1; with PA 78.9:12.6:8.5). We examined the effect that increased murine intestinal PA levels had on virulence of the AIEC strain LF82. Mice fed PA supplemented drinking water for three days prior to infection and for the duration of the infection were infected with LF82, alongside control mice that were given only sterile drinking water. Twenty-one days post infection, LF82 were recovered from the ileum and colon of infected mice. Key phenotypic features that distinguish the AIEC pathotype; adherence to and invasion of the intestinal epithelium, and the ability to form biofilms were then examined *(18)*. Adhesion to the Caco-2 human intestinal epithelial cell line by LF82 recovered from PA-fed mice was 16-fold higher than wild type LF82, while no significant change in adhesion was seen in LF82 recovered from mice not fed PA (Fig. 1b). There was no significant difference in invasion of LF82 from PA fed mice (Supplementary Fig. S2; fold change >5.25; p-value: 0.168). Examination of biofilm formation by LF82 post *in vivo* infection revealed that anaerobic biofilm formation was dramatically increased in LF82 recovered from mice fed PA relative to the wild type strain, while there was no significant difference between LF82 recovered from control non-PA fed mice and the wild type strain (Fig. 1c).

**Fig. 1.**
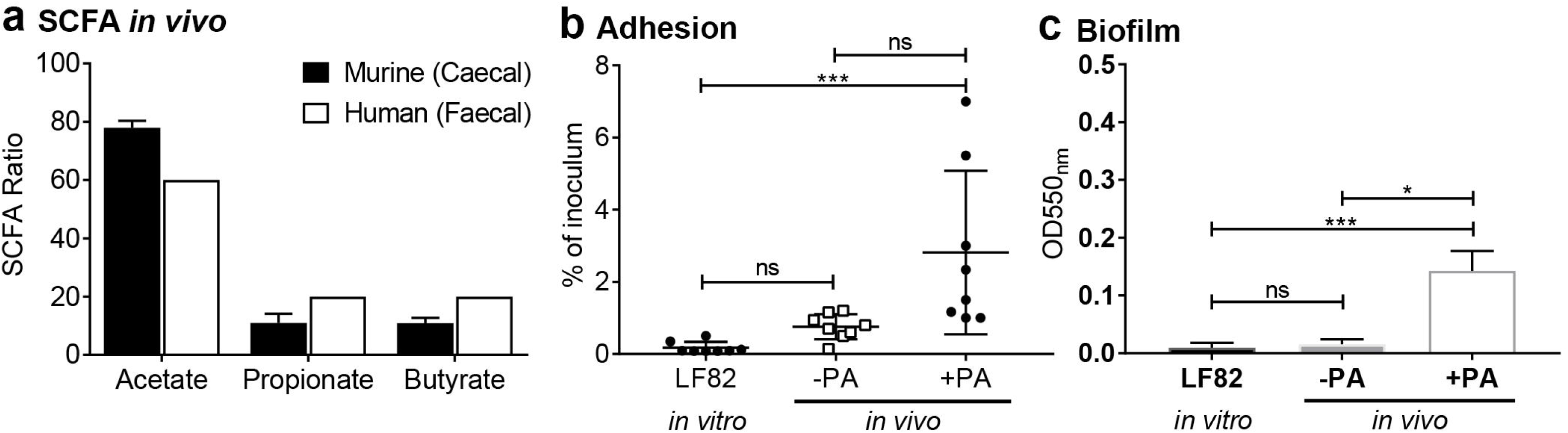
Supplementation of the murine intestine with PA raises the concentration and infection of this model with AIEC increases the virulent phenotype of the bacterium. (a) The ratio of acetate:propionate:butyrate in murine samples was calculated from the isolated caecal contents of a PBS treated mouse at 3 dpi. This was compared to published human SCFA ratios *(35)*. AIEC type strain LF82 was recovered from mice that had been given water (-PA) or water supplemented with 20 mM propionic acid (+PA). Subsequently, adhesion to Caco-2 intestinal epithelial cells (b) and ability to form biofilms in RPMI with 3% FCS (c) were assessed. *In vitro/vivo* refers to where the strains were generated. Samples were analyzed using a (a&b) multiple t-test (one unpaired t-test per row) where p<0.05 *; and a one-way ANOVA (p<0.05 *; p<0.001 ***; ns = not significant).

### The enhanced AIEC phenotype was driven by PA

We hypothesized that the enhanced virulence of LF82 was driven by the increased murine intestinal PA concentration and was independent of other factors during *in vivo* infection. To examine this, LF82 was grown in minimal media with PA as the sole carbon source (20 mM). LF82 was able to grow in PA, while a human commensal *E. coli* strain included as a control, *E. coli* F-18 *(36)*, replicated extremely poorly (Fig. 2a). Sub-culturing the bacteria over five growth cycles in PA supplemented minimal media generated a ‘PA-exposed’ strain of LF82, termed LF82-PA. LF82-PA had a significantly increased growth rate with a doubling time of 3.98 hours in PA compared to wild type LF82 at 25.59 hours (Fig. 2a). This increased growth rate was specific to PA and was not observed in nutrient rich broth (LB; Fig. 2b). These results are suprising given the well documented antimicrobial properties of PA *(20–23).*

**Fig. 2.**
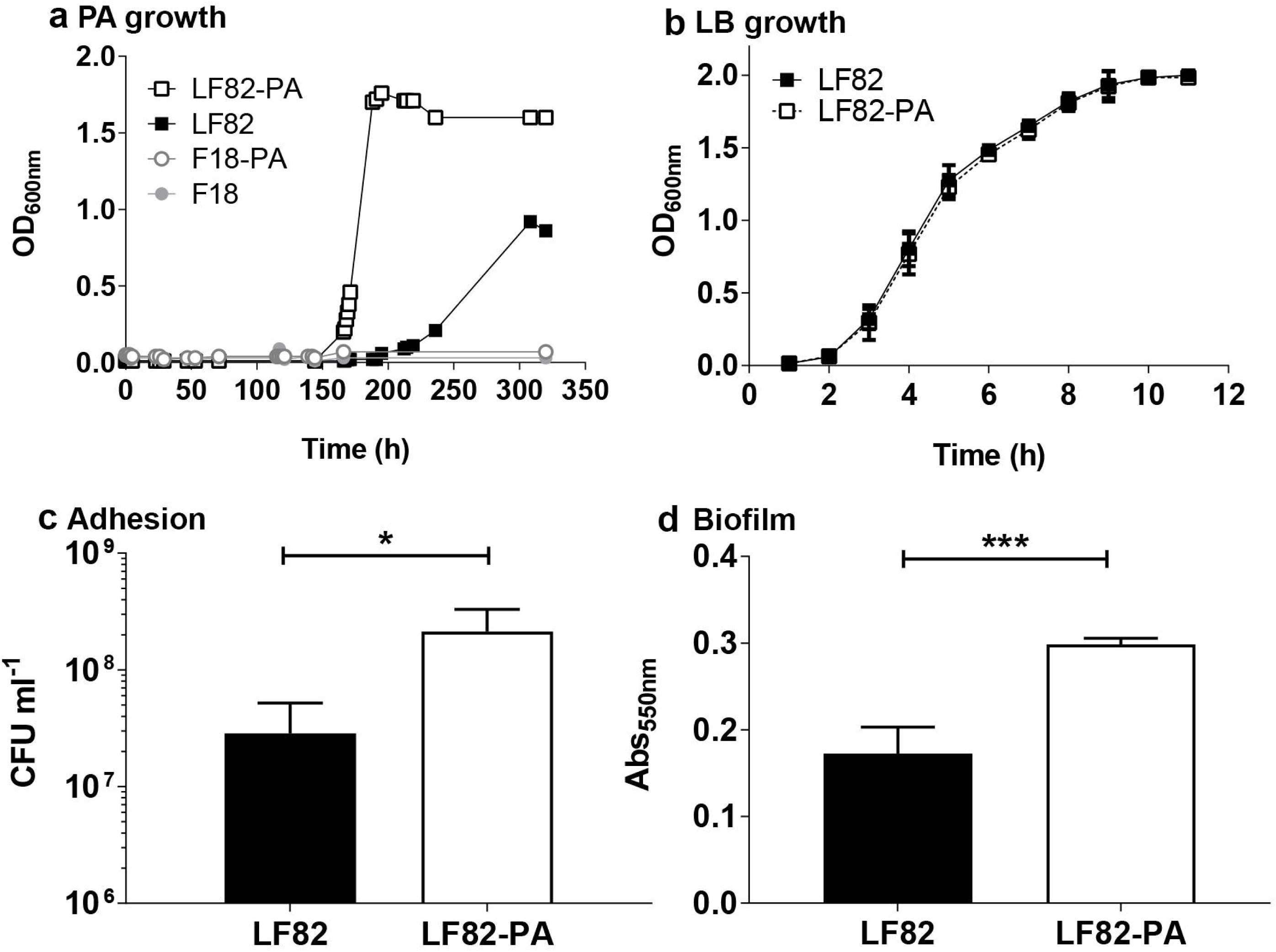
LF82 utilizes PA as a sole carbon source for growth and adaptation to PA increases virulence. (a) LF82 and commensal *E. coli* strain F-18 were grown in minimal media supplemented with 20 mM PA over a series of five successive re-cultures. Growth rate of this adapted strain, termed LF82-PA, was subsequently compared to WT LF82. (b) The growth rates of LF82 and LF82-PA were unchanged in rich LB broth. The ability of LF82 and LF82-PA strains to (c) adhere to Caco-2 intestinal epithelial cells was determined. Biofilm formation over seven days of anaerobic growth was assessed in the presence of 20 mM PA (d). Results displayed are the average of at least three biological replicates ± SD. Samples were analyzed using a (c & d) students t-test where p<0.05 *; and p<0.001 ***).

The effect on virulence of this *in vitro* exposure to PA was examined. LF82-PA showed a significant 7.45-fold increase in adherence to Caco-2 human intestinal epithelial cells when compared to LF82 (Fig. 2c), whilst similar to our strain isolated fromt the PA supplemented murine model, increased invasion by LF82-PA was observed but was not significant (Supplementary Fig. S3a, >5.26 fold; p-value = 0.3297). Anaerobic biofilm formation by LF82-PA was also significantly enhanced compared to wild type LF82 (Fig. 2d).

Direct incorporation of PA into the membrane is a mechanism employed by bacteria to minimize the toxic effects of excess PA in the environment *(37–39)*. Given that the phenotypic changes seen in AIEC such as increased adhesion, were likely to be mediated by changes in the composition of the bacterial membrane following growth on PA, we investigated this further. Gas chromatography coupled to isotope ratio mass spectrometry (GC-IRMS) using ^13^C labelled PA (1-^13^C sodium propionate) revealed that PA was not incorporated into odd chain long chain fatty acids (LCFAs). However, there was significant ^13^C-enrichment in 12 fatty acid methyl esters (FAMEs) that did not correspond to any of the 37 FAMEs in our reference standard (see *Supplementary experimental procedures*). The proximity of these labelled peaks to known LCFAs is likely indicative of incorporation of PA into methylated or branched chain fatty acids (BCFAs) as described previously during *Mycobacterium tuberculosis* growth on and detoxification of PA *(37)*. Therefore this indicated that LF82 could both metabolize and detoxify an antimicrobial that exerts potent toxic effects on a number of other intestinal pathogens *(20–23)*.

The observed changes in the bacterial membrane also rendered LF82-PA increasingly acid tolerant despite exposure in PA-supplemented minimal media being carried out at pH 7.4 (Supplementary Fig. S3b). A reduction in cell number was seen for both LF82 and LF82-PA at a pH of 3 but the LF82-PA strain survived in greater numbers for longer periods (At 20 min, LF82-PA was recovered in numbers >30.4 fold higher than LF82 [p-value = 0.0286]; at 40 min, LF82-PA was >22.7 fold higher [p-value = 0.028]). Taken collectively, these results indicate that the increased virulence observed after passage of LF82 through a PA-supplemented murine model can be replicated by exposure to PA *in vitro*.

### The PA-driven enhanced AIEC phenotype is seen in other clinical AIEC isolates

AIEC are defined phenotypically, rather than genetically, due to the lack of known distinguishing genetic determinants. Here, *E. coli* isolated from intestinal samples of paediatric patients with active CD were compared to LF82 for their ability to; adhere to and invade an intestinal epithelial cell line, replicate intracellularly in macrophages, and to form biofilms (Fig. 3). All isolates exhibited an AIEC phenotype with an ability to adhere to and invade intestinal epithelial cells, replicate within macrophages and form biofilms, both aerobically and anaerobically. There was no significant difference between the clinical isolates and the wild type LF82 strain in adherence or invasion of intestinal epithelial cells (Supplementary Table S2). While there was a reduction in intracellular replication in macrophages in strains B122 and B125, the same strains showed significantly increased biofilm formation compared to LF82. Repeated exposure of the clinical isolates to PA was undertaken as before to determine if, in a similar manner to LF82, virulence was modulated by PA. PA exposure resulted in increases in adhesion and invasion of intestinal epithelial cells and intra-macrophage replication across all isolates (Fig. 3a-c). Biofilm formation was increased under both aerobic and anaerobic growth conditions in all isolates relative to wild type (Fig. 3d-e). This data indicated that PA-induced exacerbation of the AIEC phenotype was not specific to LF82 but occurred more widely in AIEC isolated from CD patients.

**Fig. 3.**
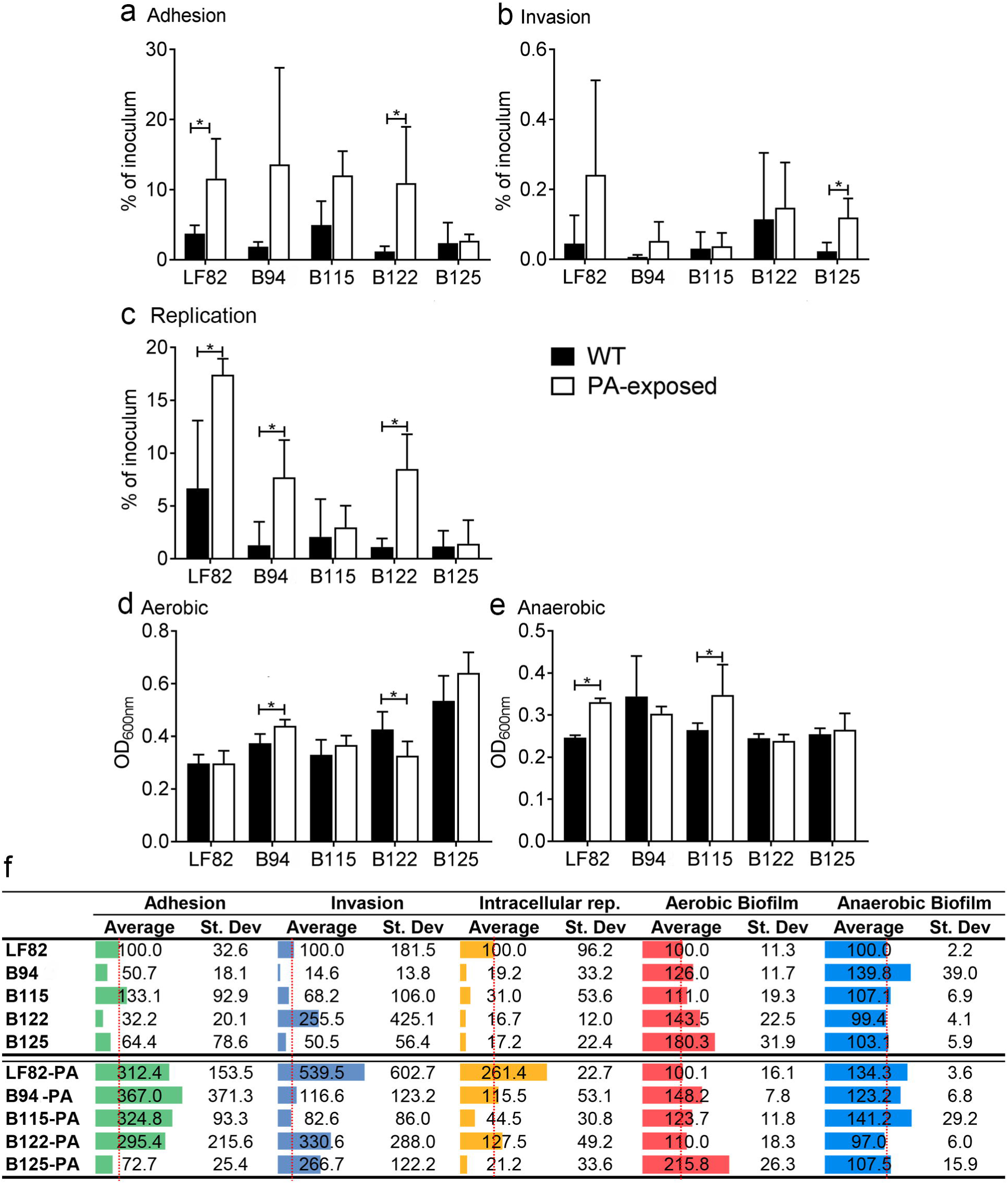
Exposure of clinical isolates to PA results in an enhanced virulence phenotype. The ability of WT and PA-exposed strains to (a) adhere to; (b) invade Caco-2 intestinal epithelial cells; (c) intracellularly replicate within macrophages; (d) form biofilms aerobically; and (e) form biofilms anaerobically, was determined. Results displayed are the average of at least two biological replicates ± SD. Samples were analyzed using two way ANOVAs (a&b) or multiple t-tests (one unpaired t-test per row; c,d&e) where p<0.05 *; and p<0.01 **). All data are summarized in (f).

### The enhanced virulence phenotype of LF82-PA is not genome-encoded and is reversible

Genome sequencing of three biological replicates of LF82-PA, exposed independently *in vitro* to PA, revealed a number of single nucleotide polymorphisms (SNPs; Supplementary File S1). However, no SNPs were conserved across all isolates. Detailed analysis of the genes and pathways in which the SNPs were identified did not lead to the identification of any candidate pathways that may explain the changes in virulence observed. However, we cannot exclude the possibility that different combinations of small genomic changes may result in the same outcome at the transcriptional level. As the virulent phenotype persists over a number of generations, and was not explained by genetic analysis, we hypothesised that the phenotype we see may be as a result of an epigenetic switch in LF82-PA. A long-term epigenetic memory switch with a role in controlling bacterial virulence bimodality was recently identified in enteropathogenic *E. coli* (EPEC) *(40)*. This ‘*resettable phenotypic switch*’ results in populations of virulent and hypervirulent genetically identical subpopulations that are retained through generations. To further examine this possibility, LF82-PA was passaged through rich (LB) media with no PA selective pressure. After five successive subcultures, this strain (LF82-PA-LB) had lost its increased growth rate in PA and its virulence phenotype reverted to that of wild type LF82 with no significant difference in adhesion of either strain to intestinal epithelial cells (Fig. 4). The epigenetic nature of this change was further confirmed through sequencing of the reverted LF82-PA-LB strains, which indicated that the SNPs present in the original LF82-PA strains were conserved, and that the changes induced by PA were not due to SNPs or mutations (Supplementary File S1). The reversion of LF82-PA-LB to wild type levels of adhesion and invasion after removal of the PA pressure was also reproducible in our clinical isolates with each reverting to their original phenotype after repeated growth in the absence of PA (Supplementary Fig. S4).

**Fig. 4.**
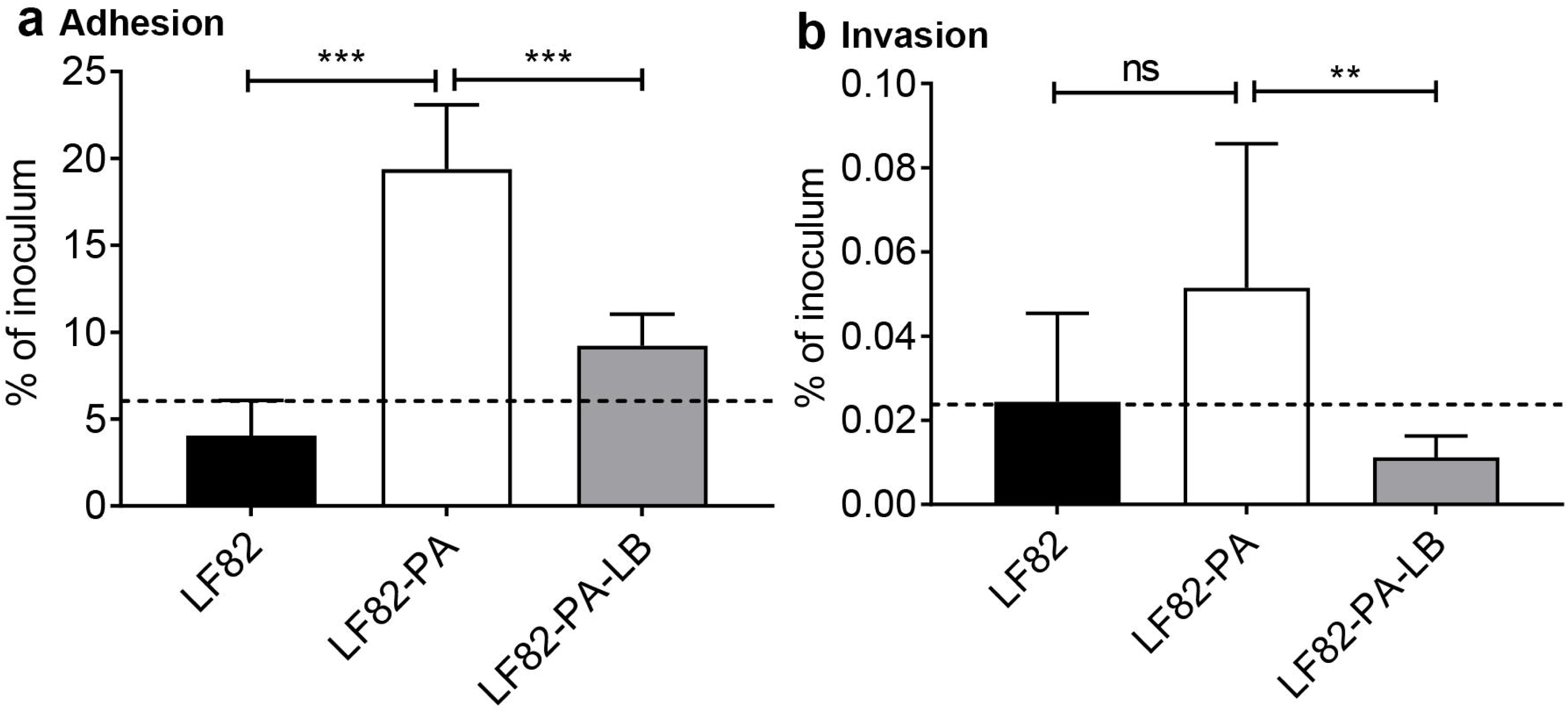
The enhanced fitness and virulence of the LF82-PA phenotype can be reversed. Continuous re-culturing (5 passages) of the LF82-PA strain in LB, generating a reverted strain (termed LF82-PA-LB) resulted in a reduction in both adherence to and invasion of Caco-2 intestinal epithelial cells. Data displayed are of two independent experimental replicates, with each experiment including three independent biological replicates. Adherence and invasion data are expressed as mean ± SD; data were analyzed using a one-way ANOVA (p<0.01 **; p<0.001 ***; ns = not significant).

### Enhanced LF82-PA virulence is driven by transcriptional changes

Given no definitive mutational basis for the observed increase in virulence was detected through genome analysis, we employed a comparative RNA-sequencing (RNA-seq) approach to probe the global transcriptional profiles of LF82 and LF82-PA grown on PA. RNA-sequencing revealed 25 differentially expressed genes (DEGs; P-value ≤0.05) between LF82 and LF82-PA (Fig. 5; Supplementary Table S3); 24 were upregulated in the LF82-PA strain and one (*mcbR;* −20.85 fold) was downregulated. Of the 25 DEGs identified by RNA-seq, 21 including *mcbR*, were validated as significantly altered by PA using qRT-PCR (Supplementary Fig. S5). Functional grouping of these 21 DEGs revealed their roles in diverse processes including biofilm formation, stress responses, metabolism, membrane integrity and transport of alternative carbon sources (Fig. 5; Supplementary Table S3). Eight DEGs have well described roles in biofilm formation further adding to our *in vitro* findings indicating that PA was a driver of adhesion and biofilm formation (Fig. 2c & d). Upregulation of another DEG, a regulator of membrane fatty acid composition *yibT*, adds further evidence for potential detoxification of PA through membrane incorporation *(37, 38)*. Therefore RNA-seq analysis indicates that PA drives changes in virulence that are fundamental to the AIEC pathotype.

**Fig. 5.**
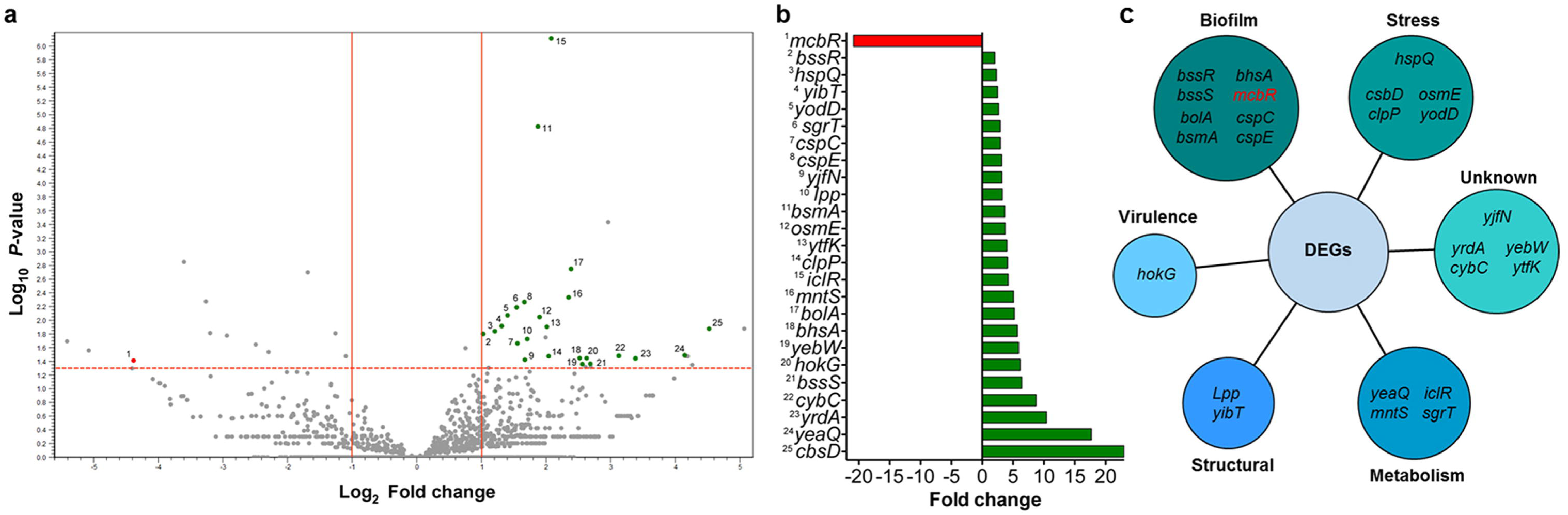
Global transcriptional shifts of LF82-PA. Volcano plot illustration of gene expression between LF82 and LF82-PA as determined by RNA-seq. Significance (Log_10_ *P*-value) and fold change cutoffs (log_2_) are indicated by the dashed and solid lines, respectively. Significantly differentially expressed genes (DEGs) are numbered and highlighted in green (upregulated in LF82-PA) and red (down regulated). Genes corresponding to ribosomal RNA coding regions were not labelled for clarity. Bar chart highlighting the fold changes in expression of the identified DEGs. Each gene is numbered and corresponds to the volcano plot. DEGs are grouped into functional categories.

### Exposure to PA in vitro significantly increases persistence of LF82 in vivo

Given our findings of PA-driven changes in virulence we determined the effect of PA exposure on long-term persistence of LF82-PA *in vivo*. Mice were again provided PA supplemented (20 mM) water for three days prior to infection, and for the 21-day duration of the infection. Mice given only sterile drinking water and infected with LF82 or LF82-PA, or treated with PBS were included as controls. In PA-fed mice, the colonization of LF82 was not significantly altered in either the ileum or colon when compared to control mice (Fig. 6). However, LF82-PA was found to persist with a greater than 20-fold increase in the ileum and a greater than 18-fold increase in the colon in these mice compared to controls (Fig. 5). No significant difference in the persistence of either strain was observed in the caecum, irrespective of the presence of PA (Supplementary Fig. S6). These data indicate that PA-exposed strains retain virulence when transferred to a new model host.

**Fig. 6.**
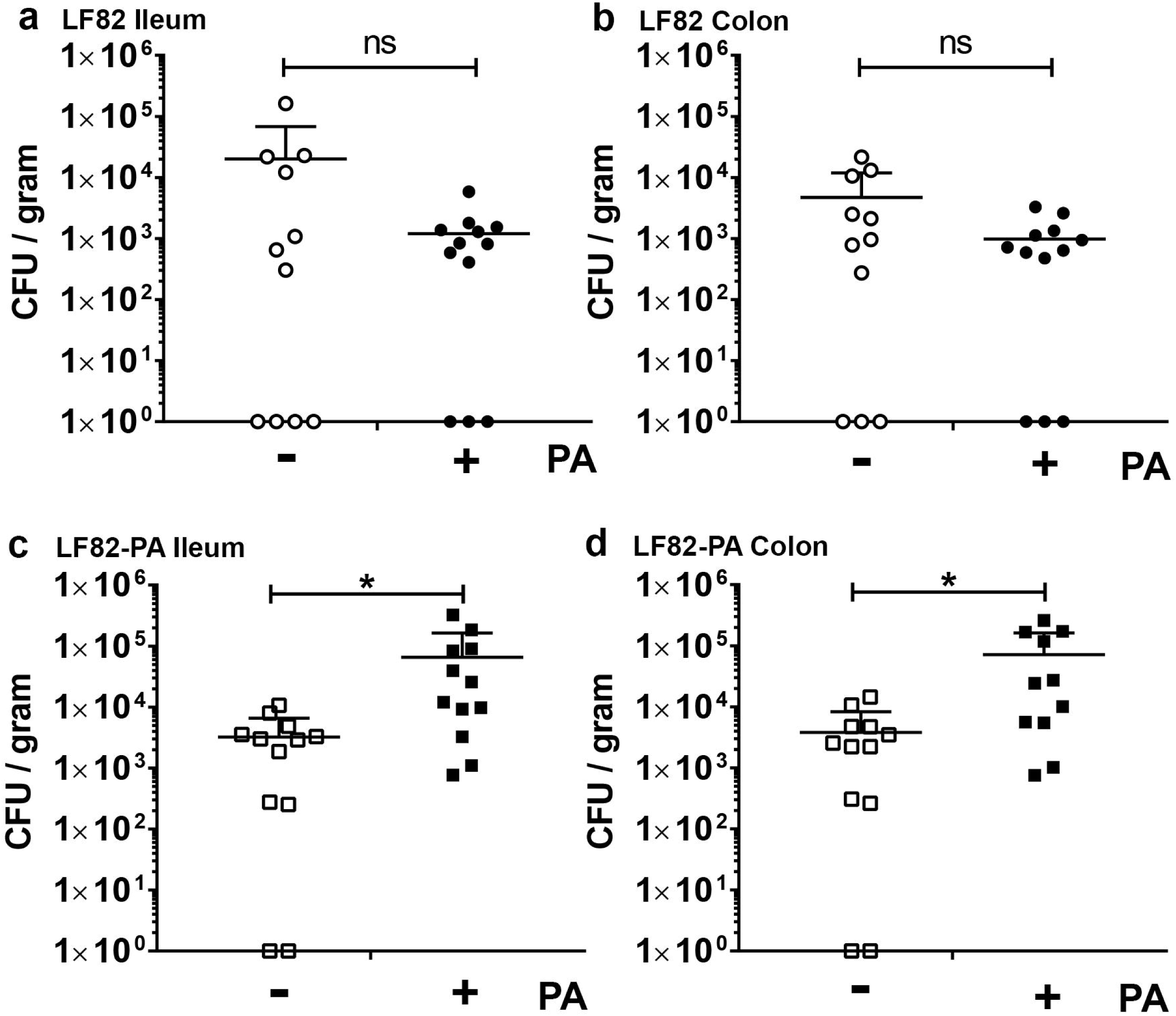
Pre-exposure of AIEC to PA combined with exogenous PA supplementation promotes colonization and long-term persistence. Drinking water was supplemented where indicated with 20mM PA and provided to male C57BL/6 mice for three days prior to infection. Persistence of LF82 (a&b) and LF82-PA (c&d) was determined in the ileum (a&c) and colon (b&d) 21 days post infection by colony counts. Data are expressed as CFU/gram of tissue ± SD and were analysed using a Students t-test; p<0.05*; ns=not significant.

## Discussion

SCFAs have significant effects on their hosts and are seen as key players in how the intestinal microbiome maintains health, modulates the immune system, controls invading pathogens, and even exerts effects on distal sites such as the brain *(41–43)*. Such positive effects have led to the suggested use of SCFAs as a therapeutic intervention strategy in inflammatory diseases, including inflammatory bowel disease *(2)*. Here however, we have shown a novel role for the SCFA PA in microbial infection, acting as a driver for virulence of the bacterial pathotype AIEC that is commonly isolated from the intestine of CD patients. Exposure of AIEC to PA, in contrast to other pathogens where it is a negative regulator of virulence, stimulated adhesion and biofilm formation and induced an upregulation of an array of genes related to virulence.

Negative regulation of bacterial virulence by PA is the basis for its widespread use in agriculture as PA clears *Salmonella* and *Campylobacter* spp. rapidly post-treatment of poultry *(20–23)*. As well as its direct toxicity to these pathogens, this inhibitory effect is mediated by negative regulation of genes critical for intestinal colonisation and is thought to be a response by these pathogens to the varying PA concentrations in the human intestine. In *Salmonella*, high concentrations of PA undermine the stability of virulence regulators such as HilD meaning *Salmonella* is less likely to colonize the lower intestine where PA levels are highest *(20)*. In direct contrast, we have shown here that PA positively regulates virulence of the AIEC strain, LF82. Increasing exposure to PA, and increasing concentrations in the murine intestine, results in a >20-fold increase in persistence as well as increasing the most notable phenotypic traits of AIEC such as adhesion to the intestinal epithelium and biofilm formation. There are limited bacteria to draw comparisons to with regards PA and virulence as few respond positively to PA given its antimicrobial properties. Enterohaemorrhagic *E. coli* (EHEC) and Citrobacter increase virulence via T3S as previously shown by Nakanishi *et al*. (2009) and Connolly *et al*. (2018), respectively. However, mycobacteria do become increasingly virulent in the presence of PA with overlapping strategies employed by mycobacteria and AIEC as both metabolize and directly incorporate PA into their membrane lipids *(38, 39, 46, 47)*. This strategy of PA incorporation in AIEC, as well as detoxifying PA, also increases resistance to pH and likely plays a role in the increased adherence noted with LF82 after PA exposure.

While PA has significant immunomodulatory properties in the intestine, disease context is highly important. While PA supplementation in certain murine models reduces disease through signalling to specific immune cells, here PA supplementation in the presence of AIEC resulted in significant overgrowth of the bacteria due to PA driven phenotypic switches that occurred *(41)*. The dramatic increase in colonization with increasing PA in the murine intestine also highlights a significant problem with murine models of intestinal disease. PA levels, along with those of other SCFAs, are significantly different in the murine intestine compared to the human intestine which is likely a contributing factor in the very different outcomes in bacterial infection in these hosts *(35)*. While differing microbiomes are a factor in susceptibility to infection, it is probable that these protective effects of the microbiome are mediated through, and dependent on, the production of SCFAs and other antimicrobial molecules by the intestinal microbiota *(42)*. Interestingly in this regard, the caecum where the majority of SCFAs are produced in the murine intestine, was distinct from the ileum and colon during infection as no significant increase in colonization by AIEC was detected here. It is possible that the levels and types of antimicrobials being produced in the caecum still proved refractory to increased AIEC colonization despite its PA-adaptation.

Our ability to recapitulate *in vitro* the effects of PA on AIEC indicates that PA in isolation exerts a significant effect on bacterial virulence. This was further demonstrated using clinical isolates of *E. coli* derived from CD patients that were exposed to PA and tested for their ability to adhere to and invade human colonic epithelial cells. While the clinical isolates were from the intestine of CD patients, it is unknown if they are true AIECs given the confusion over what constitutes the AIEC pathotype *(48)*. However, our phenotypic examination of these isolates suggests that they are likely to also be AIEC (Fig. 3). Exposure to PA induced significant increases in virulence in all four clinical strains. In comparison, a human commensal isolate F-18 was not able to adapt to PA. These data suggest that certain *E. coli* isolates recovered from the human CD intestine are readily adaptable to PA and that contrary to its effects on other pathogens PA is actually a driver for AIEC virulence and does not exert antimicrobial effects. Further analysis of the PA effect on a large range of *E. coli* pathobionts is necessary in order to definitively determine if this effect is AIEC specific.

We have shown through our work using *in vivo* models here that LF82 isolated post-PA supplementation in a murine model is more virulent than without such treatment. This indicates that rather than being directly inhibited by the antimicrobial effects of PA these strains instead show potential to be readily adaptable to the naturally higher concentrations of PA in the human intestine. Given the wide use of PA environmentally and agriculturally, it is not inconceivable that bacteria such as AIEC come into contact with such concentrations of PA, as those in animal water, feed and silage are reported to be 20 mM and higher *(20, 22, 23, 49)* Such exposure would likely make the PA concentrations in the human intestine, which increases from 1.5 mM in the ileum to 27 mM in the colon, easily tolerable to E. coli strains as we have indicated here *(20, 35)*. Horizontal transmission of strains from poultry to humans as previously seen, driven by antibiotics, would therefore seem highly possible *(28–31)*. Additionally, recent evidence has implicated the food additive trehalose as a contributory factor in the emergence and hypervirulence of two epidemic lineages of *Clostridium difficile (50)*. Therefore, while the focus rightly remains on antibiotic resistance, more work is needed to determine the long-term effects of alternative antimicrobials in generating more resistance and more virulent bacteria capable of horizontal transmission. While addressing one resistance problem we must be careful that another is not inadvertently created.

Our findings here are not without precedent. Dietary additives, and a mock Western diet, have been demonstrated to contribute to increased colonization of AIEC in murine models *(10, 11)*. This work explores the novel finding of PA as a paradoxical pro-virulence factor in AIEC, at odds with its perceived role as an antimicrobial. The growing use of PA in the Western diet, coupled with the rapid expansion of Crohn’s disease incidence in recent years, highlights the importance of this work in suggesting a potential mechanism for diet as a key driver of selection in the gut which would favour the carriage and transformation of an emerging pathogenic *E. coli* variant.

## Materials and Methods

### Bacterial strains and growth conditions

Pathogenic AIEC strain LF82 and intestinal commensal *E. coli* strain F-18 were used in this study and were cultivated on Lysogeny broth or agar. M9 minimal medium supplemented with 20 mM PA (M9-PA [20% M9 salts (32g Na_2_H_2_PO_4_2H_2_O (Merck), 12.5 g NaCl (Merck), 2.5 g NH_4_Cl (Fisher scientific), 7.5g KH_2_PO_4_ (Merck) and 400 ml H_2_O], 0.1% Trace metal solution, 0.2 mM MgSO_4_ [VWR chemicals], 0.02 mM CaCl_2_ [Fisher scientific], 1 mM Thiamine, 0.01 % 5 g/L FeCl_3_, 0.01 % 6.5 g/L EDTA, 0.1% taurocholic acid, 20 mM Sodium propionate and dH_2_O) was used for growth. Strains were grown in 100 ml of M9-PA at 37°C at 180 rpm, unless stated. Bacterial growth was measured at optical density 600nm (OD_600nm_). To obtain adapted cells, upon reaching stationary phase, cultures were back-diluted into fresh M9-PA. Strains for infection were back-diluted after overnight growth into 10 ml cultures of RPMI-1640 (Sigma) supplemented with 3% foetal calf serum (FCS) and L-glutamine. These were then grown at 37°C in a shaking incubator at 180 rpm to an OD_600nm_ of 0.6 before further dilution to give a final multiplicity of infection (MOI) of 100. Real-time PCR was conducted using bacteria grown in No-Carbon-E (NCE) media *(51)*. Twenty millimolar sodium propionate (Sigma), 1,2-propanediol (Fisher Scientific) or D-glucose (Sigma) were added with 200 nM cyano-cobalamin (Sigma) to act as an electron acceptor *(52)*. Cultures were grown overnight in LB, washed three times in NCE media with no carbon source added, and inoculated 1:100 into 10 ml NCE media containing each respective carbon source. Cultures were grown until mid-log phase (OD_600_ of 0.6) and used for RNA-extraction.

Clinical isolates (B94, B115, B122 and B125) were from the “Bacteria in Inflammatory bowel disease in Scottish Children Undergoing Investigation before Treatment” (BISCUIT) study *(53)*. Isolates B94, 115, 122 and 125 were recovered from patients with Crohn’s disease. The median (range) age was 13.7 (11.2 to 15.2), height z-score was −0.4 (−2.0 to 0.2), weight z-score was −0.7 (−3.4 to −0.1), and BMI z-score was −1.3 (−4.0 to 0.4). Symptom duration prior to diagnosis was median 7.5 months (5 to 12). 50% had granulomas present on initial histology. Phenotypes by Paris criteria *(54)* at diagnosis were: B94-colonic, non-stricturing/non-penetrating (L2, B1); B115-colonic, non-stricturing/non-penetrating (L2, B1); B122-ileocolonic, stricturing (L3, B2); B125-ileocolonic, non-stricturing/non-penetrating (L3, B1). This study is publically registered on the United Kingdom Clinical Research Network Portfolio (9633).

### Cell Culture and Maintenance

The Caco-2 human intestinal epithelial cell (IEC) line obtained from the American Type Culture Collection (ATCC) was maintained in Dulbecco’s Modified Eagle Medium (DMEM) medium (Sigma) supplemented with 10% FCS, L-glutamine and penicillin/streptomycin (Sigma). Cells were maintained at 37°C and 5% CO_2_ with regular media changes.

### Biofilm assays

Crystal violet static biofilm assays were performed as described previously *(55)*. Anaerobic culture conditions were achieved using a microaerophilic cabinet. PA was added to a final concentration of 20 mM.

### Adherence and invasion assays

Caco-2 IECs were washed once before infection and bacterial suspensions were added at an MOI of 100. The infection was allowed to proceed for 2 h at 37°C in 5% CO_2_ atmosphere. Non-adhered bacteria were washed away and the infected cells were lysed with 1% Triton X-100 for 5 min. Bacteria were serially diluted in Luria Bertani (LB) broth and spread onto LB agar plates. Total bacteria were enumerated by counting colony forming units (CFUs) after overnight incubation at 37°C. To determine bacterial invasion, cells were infected for 6 h, extracellular bacteria were then washed away and 50 µg/ml gentamycin sulphate was added for 1 h to kill any remaining cell-associated bacteria before triton X-100 treatment.

### Acid survival assays

Cultures of bacteria were grown overnight at 37°C in LB. The pH of these cultures was lowered to pH 3 using 1 M HCl. Samples were taken every 20 min for 1 h and serially diluted in LB. Dilutions were plated in triplicate onto LB agar and incubated overnight at 37°C. Colonies were counted to determine the number of surviving cells.

### Total RNA extraction and mRNA enrichment

Bacterial cultures were grown as above and mixed with two volumes of RNAprotect reagent (Qiagen, Valencia, CA, USA), before incubating for 5 min at room temperature. Total RNA was extracted, genomic DNA removed and samples enriched for mRNA as described previously by Connolly *et al.* (2016). Samples for RNA-sequencing (RNA-seq) analysis were QC tested for integrity and rRNA depletion using an Agilent Bioanalyzer 2100 (University of Glasgow, Polyomics Facility).

### Genomic analysis and SNP identification

A bacterial lawn generated from single overnight colonies of LF82 and three independent cultures of LF82-PA were resuspended in a microbank bead tube, inverted four times and incubated at room temperature for 2 min. The cryopreservative was removed and the samples sent to MicrobesNG (Birmingham University, UK) for sequencing. Genomic DNA was extracted using a Illumina Nextera XT DNA sample kit as per manufacturer’s protocol (Illumina, San Diego, USA). Samples were sequenced on the Illumina MiSeq using a 2×250 paired-end protocol, De novo assembled using SPAdes version 3.5, aligned to the reference genome using BWA-MEM 0.7.5. Variants were called using samtools 1.2 and VarScan 2.3.9 and annotated using snpEFF 4.2. Subsequent genomic analysis was performed using a combination of MAUVE, CLC genomics (Version 7.0.1), ExPASY and EMBOSS Needle. The sequence reads in this paper have been deposited in the European Nucleotide Archive under study (upload in progress).

### RNA-seq transcriptome generation and data analysis

cDNA synthesis and sequencing was performed at the University of Glasgow Polyomics Facility, essentially as described by Connolly *et al.* (2016). Briefly, sequencing was preformed using an Illumina NextSeq 500 platform obtaining 75 bp single end reads. Samples were prepared and sequenced in triplicate. Raw reads were QC checked using FastQC (Babraham Bioinformatics, Cambridge, UK) and trimmed accordingly using CLC Genomics Workbench (CLC Bio, Aarhus, Denmark). Trimmed reads were mapped to the LF82 reference genome (NCBI accession number: CU651637) allowing for 3 mismatches per read. Analysis of differential expression was performed using the Empirical analysis of DGE tool, which implements the EdgeR Bioconductor tool *(57)*. Differentially expressed genes were identified by absolute fold change (cutoffs log_2_) and a P-value of ≤ 0.05. Volcano plots were generated in CLC Genomics Workbench. The sequence reads in this paper have been deposited in the European Nucleotide Archive under study (upload in progress).

### Quantitative real-Time PCR (qRT-PCR)

cDNA was generated from total RNA using an Affinity Script cDNA Synthesis Kit (Agilent) following the manufacturer’s instructions. Levels of transcription were analyzed by qRT-PCR using PerfeCTa^®^ SYBR^®^ Green FastMix^®^ (Quanta Biosciences). Individual reactions were performed in triplicate within each of three biological replicates. The 16S rRNA and *rpoS* genes were used to normalize the results. RT-PCR reactions were carried out using the ECO Real-Time PCR System (Illumina, San Diego, CA, USA) according to the manufacturer’s specifications and the data were analyzed according to the 2^-ΔΔCT^ method *(58)*. All primers used are listed in Supplementary Table S1.

### Construction of p16S*lux*

LF82 and LF82-PA *lux* integrated strains containing the erythromycin cassette were generated using the protocol of Riedel *et al. (59)*. The bioluminescent properties of these strains allowed visualization of the establishment of infection but despite bacteria being recovered it was noted that bioluminescent signal was lost. However, upon plating the murine microbiome onto LB agar containing ampicillin (100 µg/ml), it was observed that several members of the microbiota also harboured ampicillin resistance. LB supplemented with erythromycin (500 µg/ml) did not support the growth of any microbiota species; therefore utilizing the erythromycin cassette inserted as part of the *lux* integration allowed for the selection of LF82 and LF82-PA, and was used in subsequent animal experiments.

### Animal experiments

All animal procedures were approved by an internal University of Glasgow ethics committee and were carried out in accordance with the relevant guidelines and regulations as outlined by the U.K. Home Office (PPL 70/8584). Male C57BL/6 mice aged between eight and ten weeks were obtained from The Jackson Laboratory (Envigo). Twenty millimolar sodium propionate was administered to C57BL/6 mice in drinking water three days prior to infection. Control mice were given only sterile water. Twenty-four hours prior to infection, mice were treated with an oral dose of 20 mg streptomycin before oral infection with 0.1 ml PBS (mock-infected) or with approx. 1 x 10^9^ colony forming units (CFU) of LF82 (*lux*) or LF82-PA (*lux*). Mice were euthanized 3 days after infection for colonization experiments and 21 days after infection for persistence experiments, with caecal contents collected for SCFA analysis. Ileal, caecal and colonic tissue were weighed and homogenized for enumeration of bacterial numbers. Bacterial numbers were determined by plating tenfold serial dilutions onto LB agar containing the appropriate antibiotics. After 24 h of incubation at 37 °C, colonies were counted and expressed as CFU per gram of tissue.

### SCFA analysis by gas chromatography

Faecal contents of murine caeca were isolated from PBS treated mice three dpi. The concentrations of acetate, propionate and butyrate per gram of dry weight were measured by gas chromatography as previously described *(60)* and expressed as a ratio for comparison to the known human acetate:propionate:butyrate SCFA ratio *(35)*. The effect of PA supplementation in drinking water on PA levels was measured by extraction of caecal contents from PA treated and untreated mice and the levels of SCFAs again calculated per gram of dry weight.

### Statistical analysis

Values are represented as means and standard deviation. All statistical tests were performed with GraphPad Prism software, version 7.0c. All replicates in this study were biological; that is, repeat experiments were performed with freshly grown bacterial cultures, immortalized cells and additional mice, as appropriate. Technical replicates of individual biological replicates were also conducted, and averaged. Significance was determined using t-tests (multiple and individual as indicated in the figure legends) and one-way ANOVA corrected for multiple comparisons with a Tukey post hoc test (as indicated in the figure legends). RT-PCR data was log-transformed before statistical analysis. Values were considered statistically significant when *P* values were \**P* < 0.05, \*\**P* < 0.01, \*\*\**P* < 0.001.

## Supporting information

Supplementary Experimental Procedures

## General

The LF82 strain was a kind gift from Professor Daniel Walker, University of Glasgow.

## Author contributions

MJO designed and performed the experiments, analysed the data and prepared the manuscript. SAJ assisted in experimental design, animal experiments and RT-PCR. LMM, RJG and DGES assisted in experimental design. AM provided technical assistance throughout the experiments. JPRC and AJR advised on and assisted in RNA-seq analysis. CCM generated the primary LF82-PA strain. HEH and HMW assisted in animal experiments. EF was involved in developing the initial concept for the study. KG advised on and conducted SCFA experiments. DM conducted GC-IRMS experiments. GH & RH provided BISCUIT isolates and relative clinical information. DMW was awarded the funding, developed the initial concept, designed experiments and prepared the manuscript. All authors contributed to editing the manuscript for publication.

## Supplementary Figure Legends

Supplementary Fig. S1 The murine and human SCFA ratios are not comparable; Supplementary Fig. S2 *In vivo* exposure of AIEC to PA increases *in vitro* invasion of intestinal epithelial cells; Supplementary Fig. S3 LF82 adaptation to PA shows slightly increased invasion and displays an acid tolerant phenotype; Supplementary Fig. S4 The enhanced PA-phenotype is reversible in *E. coli* isolates recovered from clinical samples. Supplementary Fig. S5 qRT-PCR validation of transcriptional changes of LF82-PA; Supplementary Fig. S6 Exogenous PA supplementation does not alter colonization and long-term persistence of AIEC in the murine caecum.

