## Supplementary Information for "Propionic acid promotes the virulent phenotype of Crohn’s disease-associated adherent-invasive *Escherichia coli*"

### Title


**Supplementary Figure Legends:** Supplementary Fig. S1 The murine and human SCFA ratios are not comparable; Supplementary Fig. S2 *In vivo* exposure of AIEC to PA increases *in vitro* invasion of intestinal epithelial cells; Supplementary Fig. S3 LF82 adaptation to PA shows slightly increased invasion and displays an acid tolerant phenotype; Supplementary Fig. S4 The enhanced PA-phenotype is reversible in E. coli isolates recovered from clinical samples; Supplementary Fig. S5 qRT-PCR validation of transcriptional changes of LF82-PA; Supplementary Fig. S6 Exogenous PA supplementation does not alter colonization and long-term persistence of AIEC in the murine caecum.

**Supplementary Figure Legends:**

**Supplementary Fig. S1The murine and human SCFA levels are not comparable.** Levels of murine caecal SCFAs were determined by dry weight after supplementation of the drinking water with 20mM PA.

**Supplementary Fig. S2 *In vivo* exposure of AIEC to PA increases *ex vitro* invasion of intestinal epithelial cells.** Invasion of caco-2 intestinal epithelial cells was assessed. *In vitro/vivo* refers to where the strains were generated. Samples were analyzed using a one-way ANOVA (ns = not significant).

**Supplementary Fig. S3 LF82 adaptation to PA shows slightly increased invasion and displays an acid tolerant phenotype.** The ability of LF82 and LF82-PA strains to (a) invade Caco-2 intestinal epithelial cells was determined. (b) Tolerance of LF82 and LF82-PA to acidic conditions was monitored at pH 3.0. Results displayed are the average of at least three biological replicates ± SD. Samples were analyzed using a students t-test where $p<0.05$ *; and ns is not significant.

**Supplementary Fig. S4 The enhanced PA-phenotype is reversible in *E. coli* isolates recovered from clinical samples.** The ability of clinical isolates to adhere to (a) and invade (b) Caco-2 intestinal epithelial cells was compared to LF82 before exposure to PA, after exposure to PA, and after removal of the PA pressure (through subculture in rich media). Results displayed are the average of at least two biological replicates ± SD. Samples were analyzed using a students t-test where $p<0.05$ *; and ns is not significant.

**Supplementary Fig. S5 qRT-PCR validation of transcriptional changes of LF82-PA.** qRT-PCR was conducted on LF82-PA grown in minimal media supplemented with 20 mM PA. Relative fold change was measured against LF82 grown under the same conditions, using *16S* as a control. Three independent biological replicates were performed. Data are expressed as relative fold change ± SD and were analyzed using a one-way ANOVA with Tukey; $p<0.05$ *.

**Supplementary Fig. S6 Exogenous PA supplementation does not alter colonization and long-term persistence of AIEC in the murine caecum.** Drinking water was supplemented where indicated with 20mM PA and provided to male C57BL/6 mice for three days prior to infection. Persistence was determined 21 days post infection by colony counts. Data are expressed as CFU/gram of tissue ± SD and were analysed using a Students t-test; ns=not significant.
