## Supplementary Experimental Procedures for "Propionic acid promotes the virulent phenotype of Crohn’s disease-associated adherent-invasive *Escherichia coli*"

***Measurement of 1-$^{13}$C-PA incorporation by gas chromatography coupled to isotope ratio mass spectrometry (GC-C-IRMS).*** LF82-PA was grown as before in M9 minimal medium here supplemented with 20 mM 1-$^{13}$C-PA. Cultures were harvested at OD 0.6, pelleted and washed with PBS. Air-dried cells were treated using a saponification and methylation procedure to produce fatty acid methyl esters (FAMEs) of all cell fatty acids. Briefly, to air dried cells was added 1 ml of methanol:heptane:toluene:2,2-dimethoxypropane:conc $H_2SO_4$ (39:34:20:5:2 by vol) and samples vortexed and then heated at 80°C for 30 mins. Upon cooling, 100µl of the upper heptane phase containing FAMEs was extracted to a clean vial ready for analysis. Samples were analyzed using gas chromatography coupled to isotope ratio mass spectrometry through a combustion interface (GC-C-IRMS). FAMES separated by GC (Agilent 6890, ZB-FFAP column (30 m x 0.25 mm x 0.25 µm), He carrier (2ml/min), temperature programme of 80°C start followed by 7.5°C / min to 150°C, 2°C / min to 225°C and finally 5 min dwell at 225°C) and eluting FAMEs were oxidized to $CO_2$ over hot copper oxide (GVI Isochrome, Manchester, UK) in a He flow to the IRMS. An open split design allowed a portion of the eluting CO2 in He to enter the IRMS where ions at mass to charge (*m/z*) 44, 45 and 46 were analyzed continuously and identified peaks were integrated against a reference $CO_2$ peak to yield the background and Craig corrected $^{13}C/^{12}C$ ratio expressed in the normal units δ13C (per mil) versus the internationally accepted scale for $^{13}C/^{12}C$ measurements, VPDB. Samples were bracketed by a certified reference FAME mix (Supelco® 37 Component FAME Mix, Sigma-Aldrich, UK; containing Butyate, Hexanoate, Octanoate, decanoate, Undecanoate, Laurate, tridecanoate, tetradecanoate, Myristoleic, Pentadecanoate, Cis-10-pentadecanoic, Palmitate, palmitoleic, heptadecanoic, cis-10-Heptadecenoic, octadecanoic, trans-9-Elaidic, cis-9-Oleic, Linolelaidic, linoleate, Arachidate, gamma-Linolenic, cis-11-eicosenoate, Linolenate, heneicosanoate, cis-11,14-Eicosadienoic, docosanoate, cis-8,11,14-Eicosatrienoic, Erucate, cis-11,14,17-Eicosatrienoic, tricosanoate, cis-5,8,11,14-Eicosatetraenoic, cis-13-16-Docosadienoic, lignocerate, cis-5,8,11,14,17-Eicosapentaenoate, Nervonate, cis-4,7,10,13,16,19-Docosahexaenoate) to retention time lock for 37 odd and even chain FAMEs.
