## Supplementary figures and images for "Propionic acid promotes the virulent phenotype of Crohn’s disease-associated adherent-invasive *Escherichia coli*"

### Supplemental Data 1

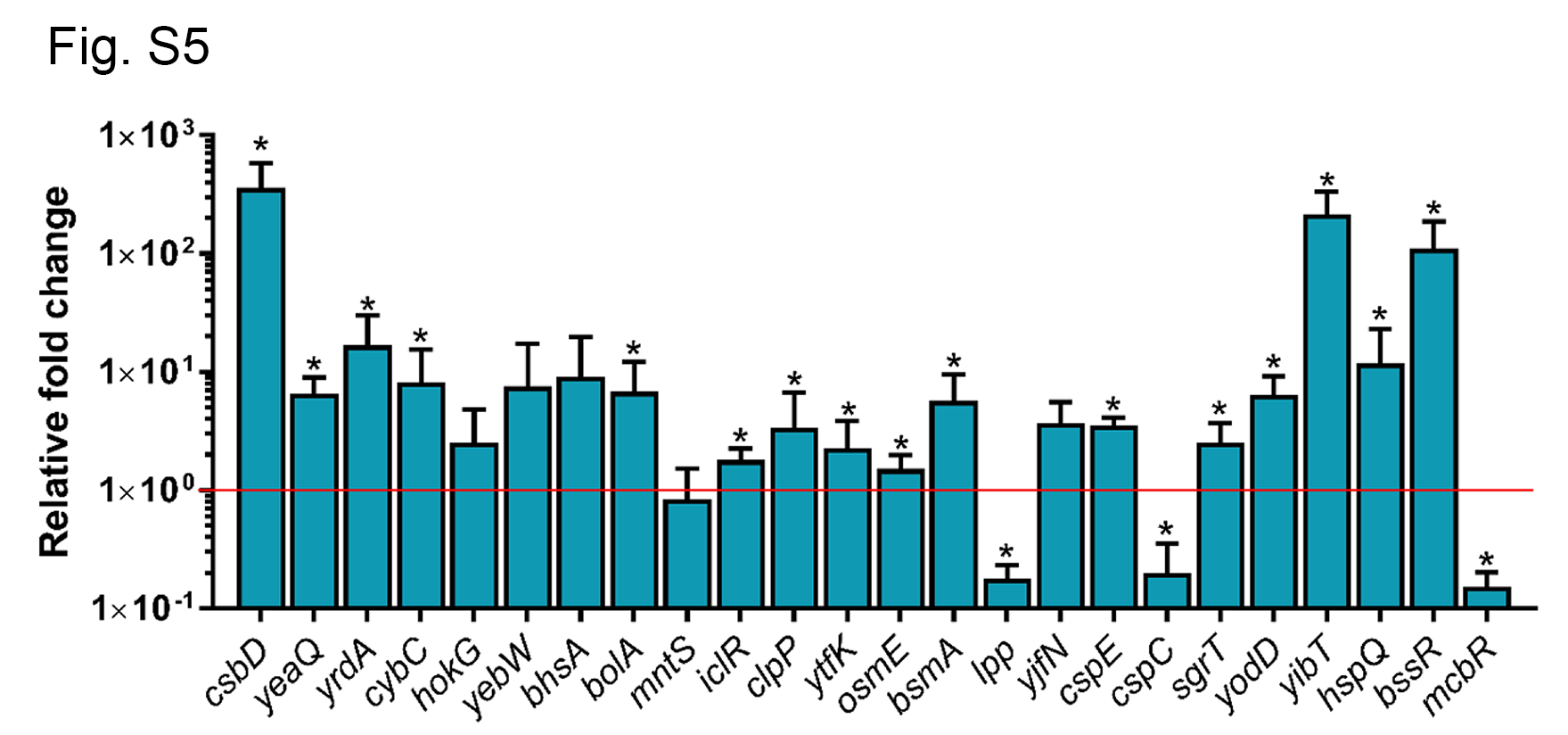

### Supplemental Data 2

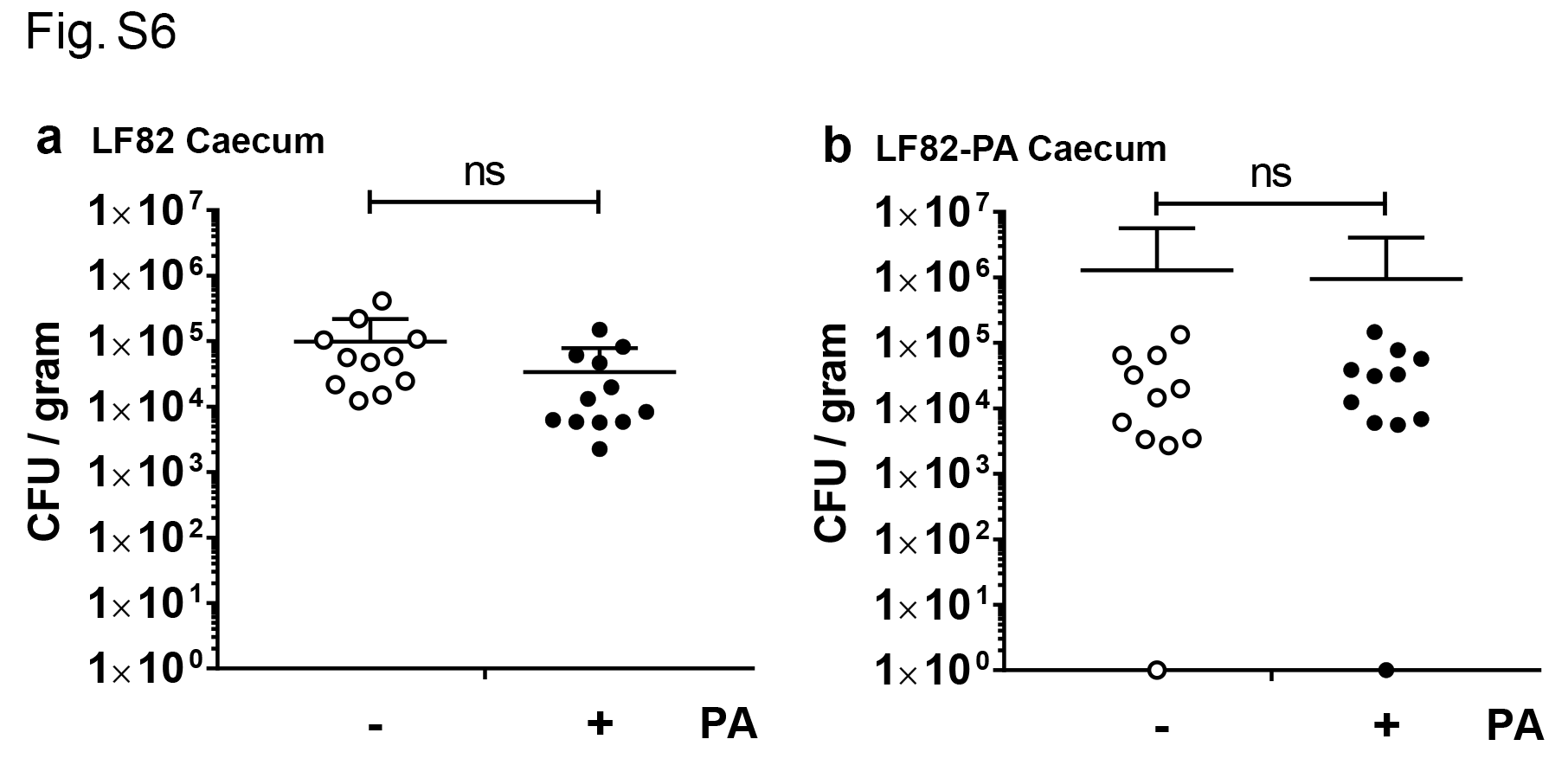

### Supplemental Data 3

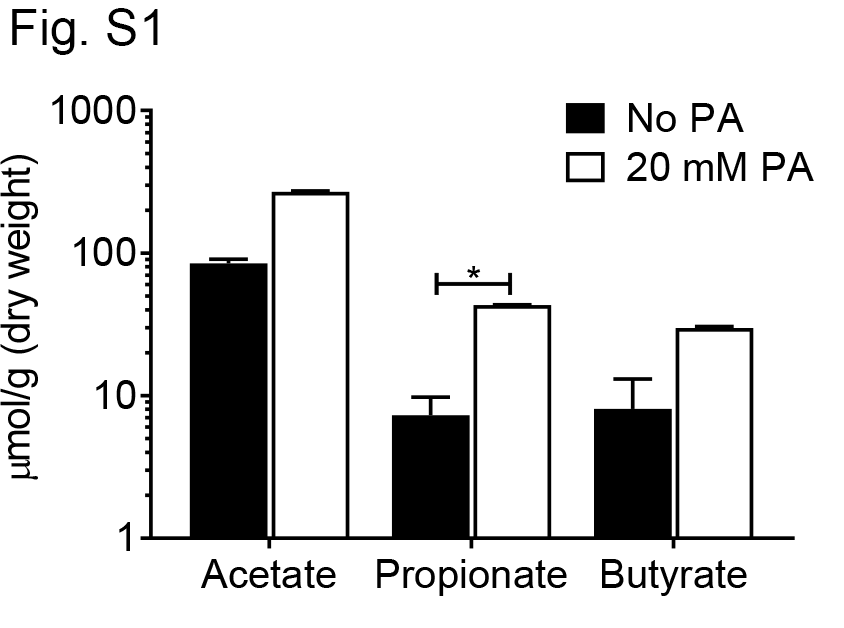

### Supplemental Data 4

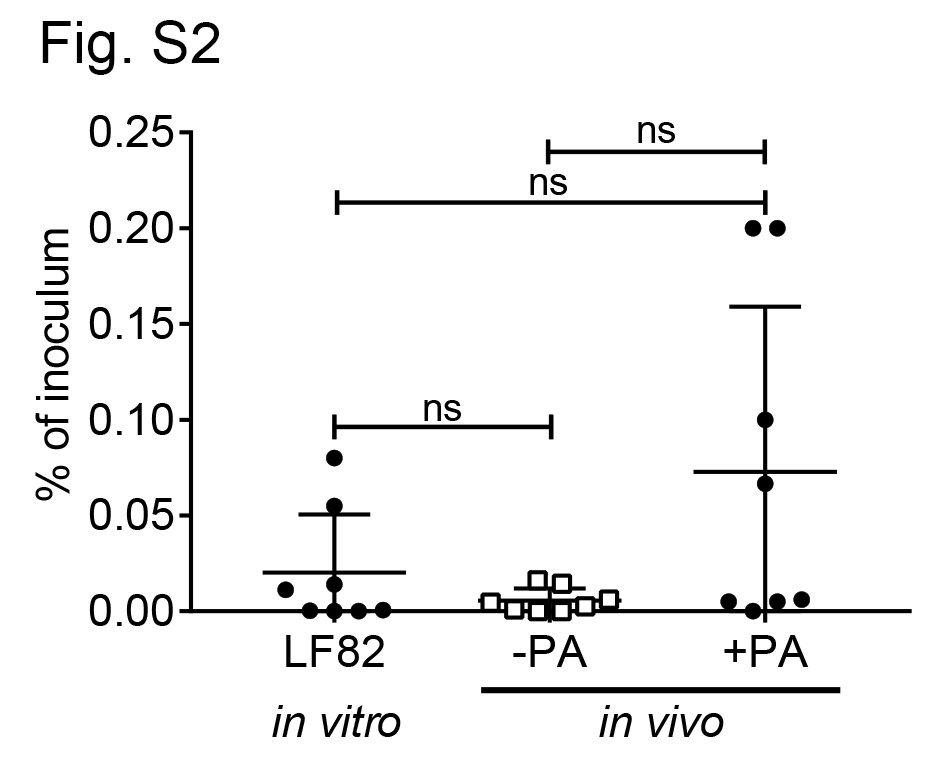

### Supplemental Data 5

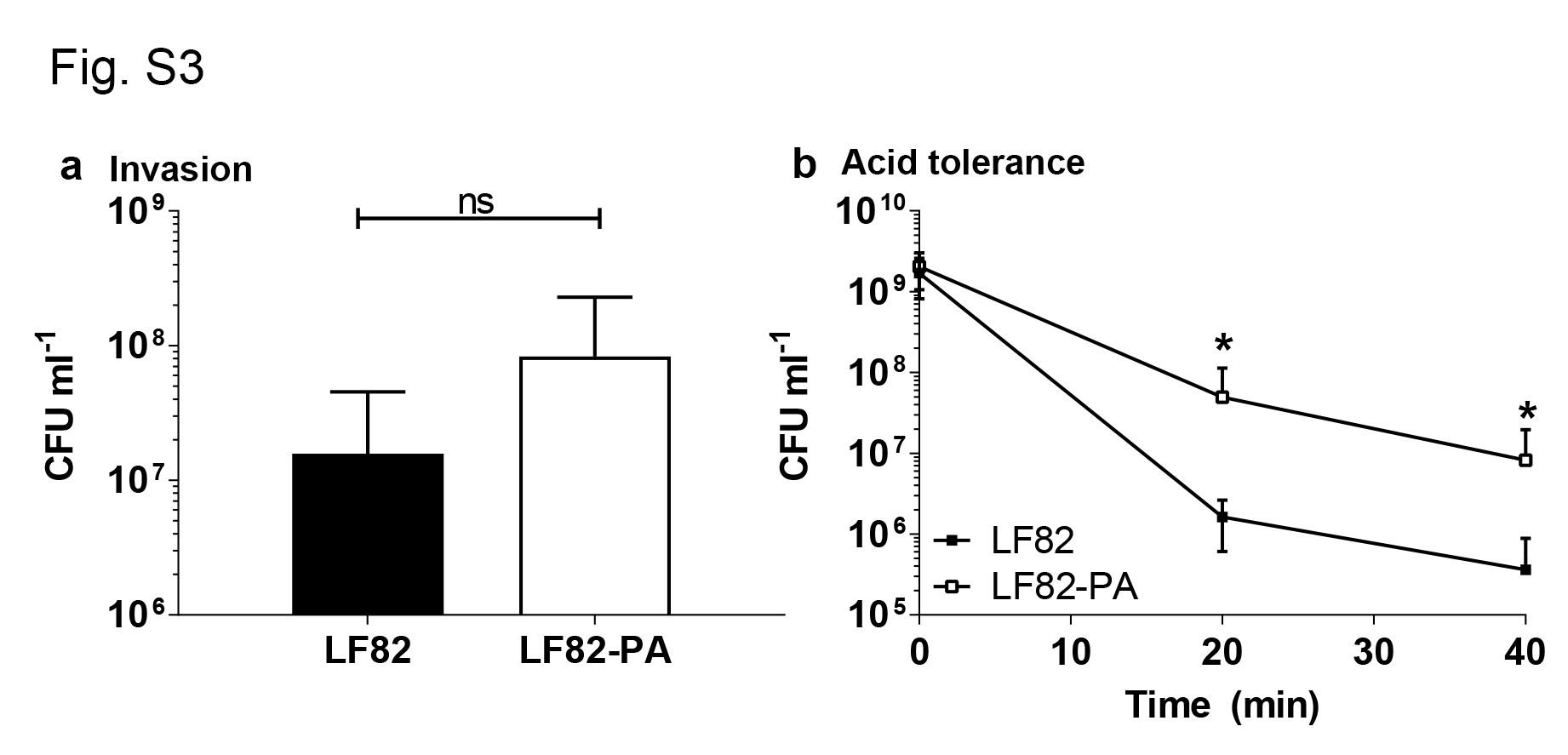

### Supplementary Fig. S4

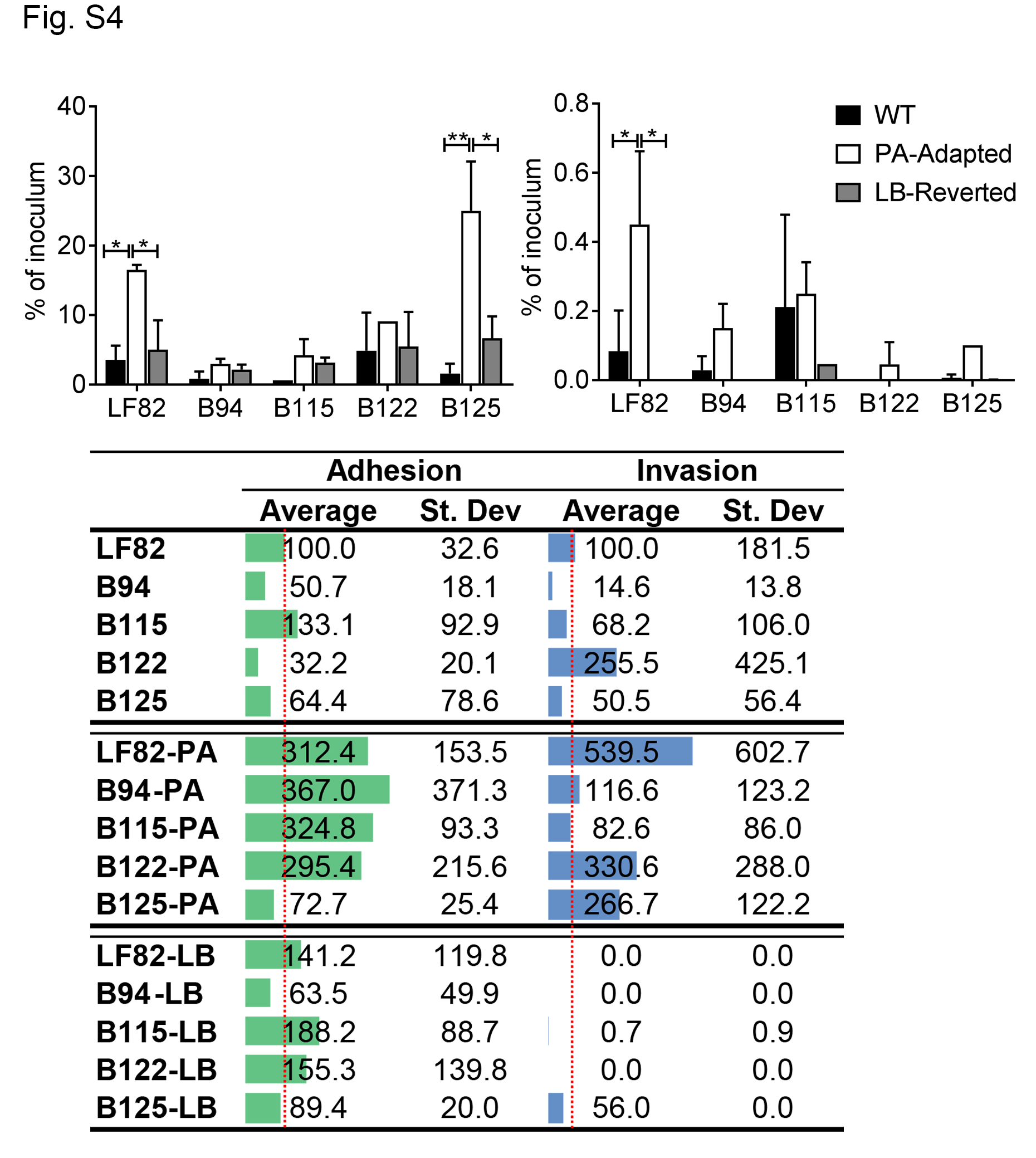
